# The C-type lectin domain of CD62P (P-selectin) is an integrin ligand

**DOI:** 10.1101/2022.08.17.504309

**Authors:** Yoko K Takada, Yoshikazu Takada

## Abstract

CD62P (P-selectin) is confined to the inside of platelets and endothelial cells, and is translocated to the surface upon activation of platelets or endothelial cells. In current models, CD62P recognizes sialyl-Lewis X on PSGL-1 and mediates rapid rolling of leukocyte over vascular surfaces during the initial steps in inflammation. Docking simulation using integrin αvβ3 as a target predicted that the C-type lectin domain of CD62P is a potential integrin ligand. It has not been tested if CD62P binds to integrins. Here we describe that the lectin domain of CD62P specifically bound to soluble integrins αvβ3, αIIbβ3, α4β1 and α5β1. Known inhibitors of CD62P-PSGL-1 interaction did not suppress the binding of the lectin domain to integrins. We found that the R16E/K17E mutation in the predicted integrin-binding interface of the lectin domain strongly inhibited CD62P binding to αIIbβ3 and αvβ3 in 1 mM Mn^2+^. R16E/K17E is outside of the glycan binding site. Mutating Glu-88 to Asp (the E88D mutation) in the lectin domain, which is known to strongly disrupt glycan binding, only slightly affected integrin binding, indicating that glycan binding and integrin binding sites are distinct. Also, the lectin domain of CD62P supported cell adhesion in a cation-dependent manner. CD62P-integrin interaction is potentially important since integrins are widely expressed compared to PSGL-1, which is limited to leukocytes. These findings indicate that CD62P-integrin interaction plays potentially important role in a wide variety of cell-cell interaction in addition to CD62P-glycan interaction.

## Introduction

CD62P (CD62P), a member of the selectin family, has been identified as a Ca^2+^-dependent receptor for myeloid cells that binds to carbohydrates on neutrophils and monocytes. CD62P is stored in the α-granules of platelets and Weibel-Palade bodies of endothelial cells (1). CD62P is transferred to the surface upon activation of platelets (2) or endothelial cells (3). The extracellular region of CD62P is composed of three different domains like other selectin types; a C-type lectin-like domain in the N-terminus, an EGF-like domain and a complement-binding protein-like domains having short consensus repeats (~60 amino acids). CD62P is anchored in transmembrane region followed by a short cytoplasmic tail region (4). CD62P is known to recognize sialyl-Lewis X and mediate rapid rolling of leukocytes over vascular surfaces during the initial steps in inflammation through interaction with CD62P glycoprotein ligand-1 (PSGL-1) (5).

CD62P is a major therapeutic target for cardiovascular diseases, inflammation and cancer metastasis (6). However, CD62P binding to sialyl-Lewis X has exclusively been targeted for drug development.

Integrins are a family of cell-surface αβ receptor heterodimers that bind to extracellular matrix ligands (e.g., fibronectin, fibrinogen, and collagen), cell-surface ligands (e.g., ICAM-1 and VCAM-1), and soluble ligands (e.g., growth factors) (7). By virtual screening of protein data bank (PDB) with integrin headpiece as a target using docking simulation, we have discovered many new integrin ligands.–These ligands are shown to bind to the classical ligand-binding site of integrins (site 1). We also found that several ligands, including CX3CL1 (8) and sPLA2-IIA (9), bind to the allosteric binding site of integrins (site 2), which is on the opposite side of site 1 in the integrin headpiece, and allosterically activated integrins.

Here we describe that we identified the C-type lectin domain of CD62P as a potential integrin ligand by virtual screening of protein data bank. Since the Pubmed search did not find a report that described that integrins interact with CD62P, we decided to pursue possible integrin-CD62P interaction. In CD62P-αvβ3 docking model, CD62P is predicted to bind to the classical ligand-binding site of integrins (site 1). The integrin-binding site in CD62P is predicted to be distinct from that of glycan-binding site. We identified amino acid residues critical for integrin binding in the lectin domain by introducing mutations in the predicted integrin-binding site (e.g., the K16E/R17E mutation). The E88D mutation that is known to block glycan binding (10) minimally affected integrin binding. Therefore, we propose that CD62P acts as an integrin ligand on activated endothelial cells or on activated platelets, and that CD62P mediates cell-cell interaction by binding to integrins, in addition to mediating glycan binding and rolling. In addition, the lectin domain of CD62P bound to site 2 and activated integrins. The present study has biological significance since integrins are widely expressed compared to PSGL-1, which is limited to leukocytes. Integrin-CD62P interaction will mediate cell-cell interaction between different cell types, including platelets, endothelial cells, leukocytes, and cancer cells.

## Results

### The lectin domain of CD62P specifically binds to soluble integrins αvβ3 and αIIbβ3

We virtually screened the protein data bank (PDB) for potential integrin ligands using docking simulation using integrin αvβ3 (1L5G.pdb, open headpiece) as a target. The simulation predicted that the C-type lectin domain of CD62P as a potential integrin ligand. This prediction is not consistent with current models of CD62P, which recognizes sialyl-Lewis X and mediates rapid rolling of leukocyte over vascular surfaces during the initial steps in inflammation by binding to PSGL-1.

To address this prediction we studied if the lectin domain directly binds to integrins.

We used the C-type lectin domain (residues 1-117) and the combined lectin and EGF-like domain (residues 1-158) (Fig. 1a). We found that soluble αvβ3 and αIIbβ3 bound to the lectin domain well in ELISA-type binding assays in 1 mM Mn^2+^ (TH-1 mM Mn^2+^) in a dosedependent manner (Fig. 1b). The lectin domain showed stronger (approx. 2x) binding to integrins than the combined lectin and EGF-like domains (Fig. 1c), indicating that the lectin domain is primarily involved in integrin binding.

**Fig. 1.**
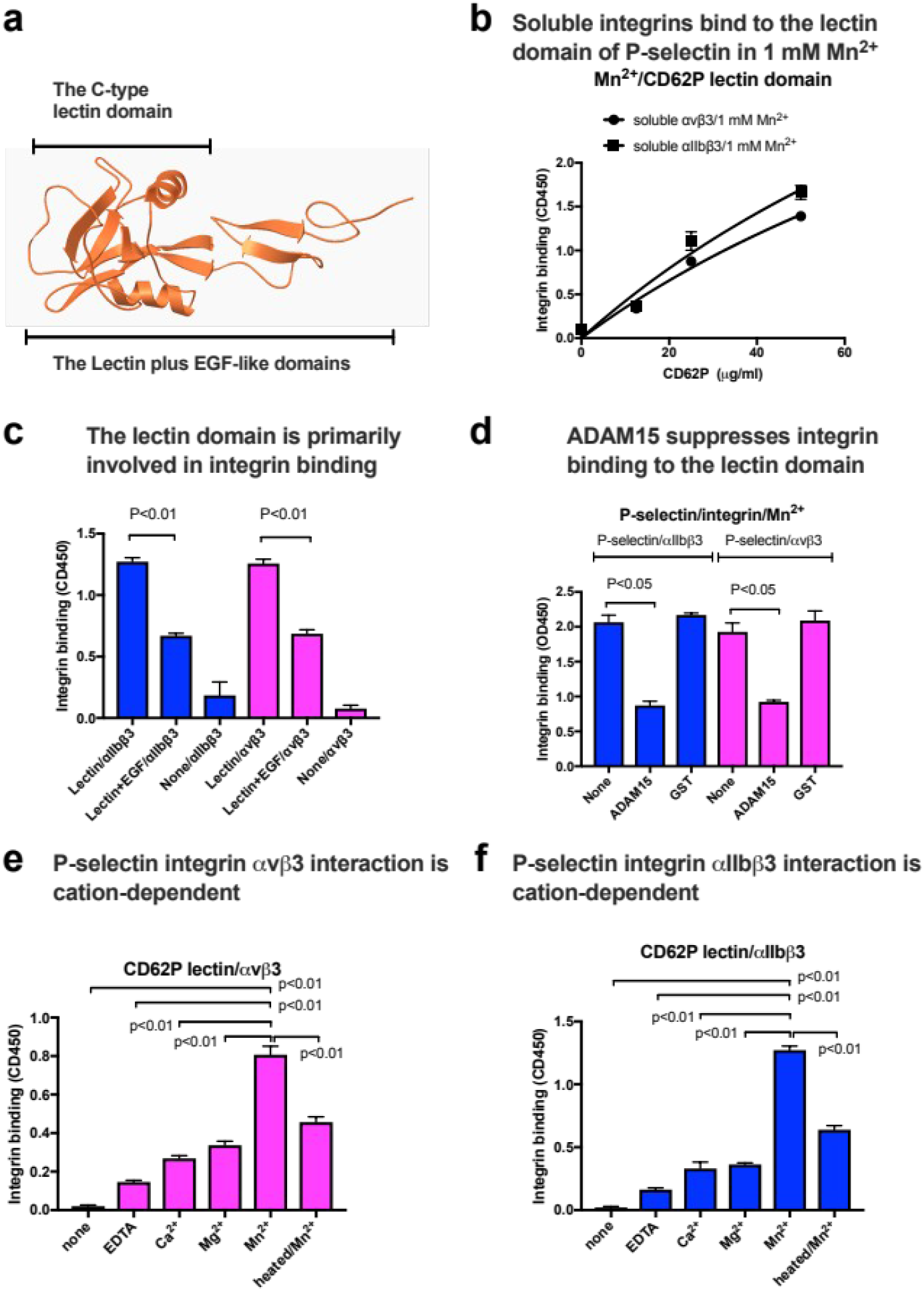
The lectin domain bound to soluble integrins αvβ3 or αIIbβ3 in ELISA-type binding assays. <u>a. The C-type lectin and the EGF domains in the crystal structure of CD62P. b. Binding of soluble integrins to immobilized lectin domain of CD62P.</u> Wells of 96-well microtiter plate were coated with the lectin domain of CD62P and the remaining protein binding sites were blocked with BSA. Wells were incubated with soluble integrin αvβ3 or αIIbβ3 (1 μg/ml) for 1 hr in 1 mM Mn^2+^ and bound integrins were quantified using anti-β3 mAb and anti-mouse IgG conjugated with HRP. <u>c. The C-type lectin domain binds better to soluble integrins than the combined lectin and EGF domains (at coating conc. of 50 μg/ml). d. The disintegrin domain of ADAM15, another ligand for αvβ3 or αIIbβ3 suppress binding of soluble integrins to the lectin domain. e. and f. The binding of soluble integrins to immobilized CD62P lectin domain (coating concentration at 50 μg/ml) in 1 mM different cations.</u> Data is shown as means +/- SD (n=3). Statistical analysis was performed by ANOVA in Prism 7.

These findings suggest that the lectin domain of CD62P is a ligand for αIIbβ3 and αvβ3. We focused on the lectin domain-integrin interaction throughout the study. However, cRGDfV or 7E3 (anti-β3) did not affect the binding of the lectin domain to soluble αvβ3. We thus studied if known ligand for integrins αIIbβ3 (11) and αvβ3 (12) competes for binding to these integrins. The ADAM15 disintegrin domain has been reported to be a specific ligand for αvβ3 and αIIbβ3. We found that ADAM15 disintegrin fused to GST suppressed the binding of soluble αvβ3 or αIIbβ3 to immobilized lectin domain (Fig. 1d), indicating that the lectin domain competes with ADAM15 disintegrin for binding to integrins. Therefore, the CD62P lectin domain is a specific ligand for αvβ3 and αIIbβ3.

Heat-treatment reduced integrin binding, indicating that the lectin domain have to be properly folded for integrin binding (Fig. 1e and 1f). Also, the lectin domain showed cation-dependency for binding to integrins αvβ3 and αIIbβ3 (1 mM Mn^2+^>Mg^2+^> Ca^2+^>EDTA), which is similar to that of known integrin ligands. These findings are consistent with the idea that the lectin domain is an integrin ligand.

### The integrin-binding site and glycan-binding site are distinct

We generated a model of interaction of integrin, CD62P, and PSGL-1 by superposing the docking model and the PSGL-1-CD62P complex. The model predicts that PSGL-1 peptide (605YEYLDYDFLPETEP618) in the PSGL-1-CD62P lectin domain complex (1g1s.pdb) (13) bind to CD62P to integrin αvβ3 without steric hindrance (Fig. 2a). We selected several amino acid residues in the integrin binding interface of CD62P (Arg16/Lys17, Lys58, Lys66/Lys67, Lys84/Arg85) for mutagenesis to Glu (Fig. 2b).

**Fig. 2.**
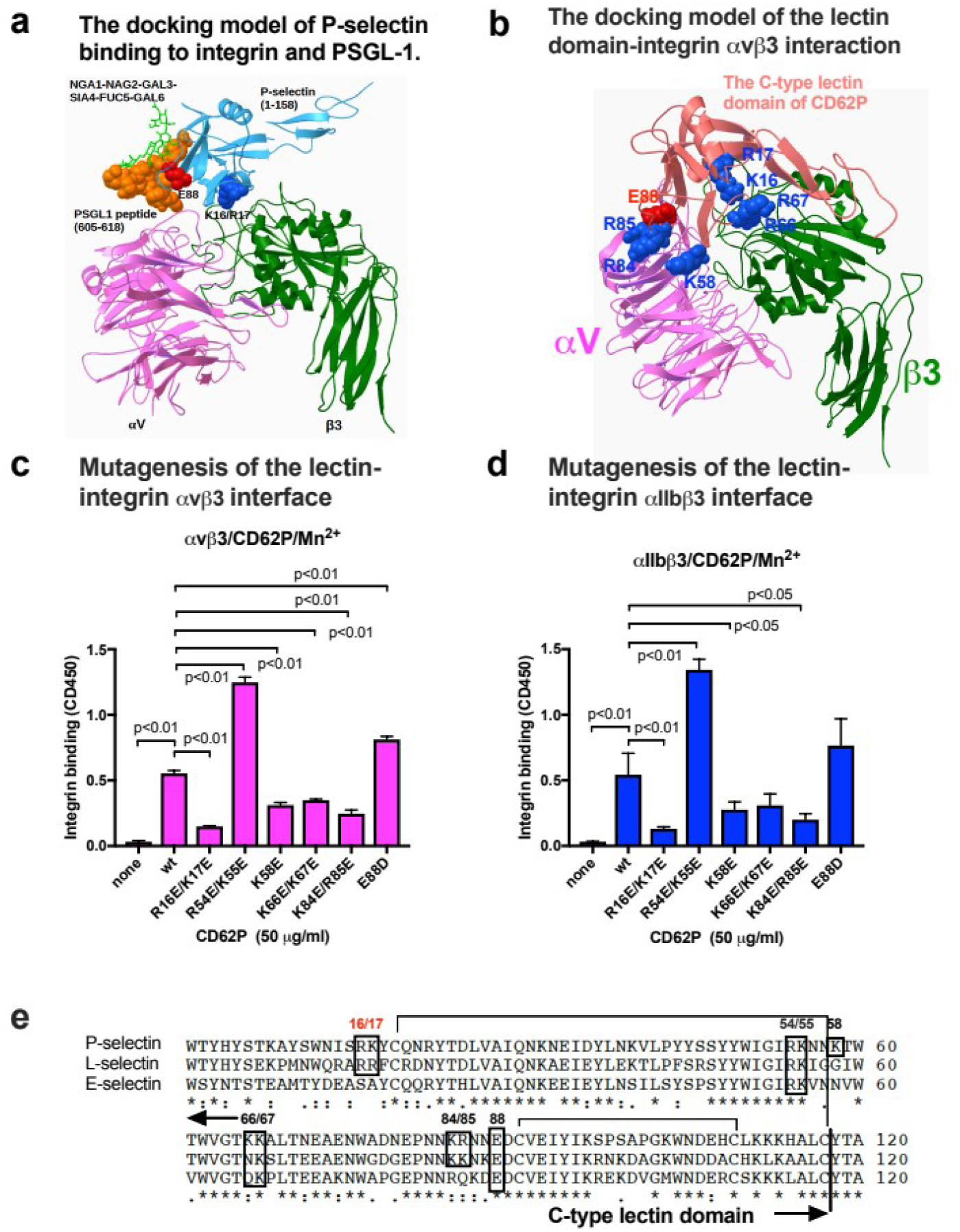
<u>CD62P binding to integrin and PSGL-1.</u>. <u>a. The CD62P-αvβ3 docking model was superposed with the crystal structure of CD62P-PSGL1 peptide complex (1g1s.pdb).</u> The superposed model predicts that integrin-binding site and PSGL-1-binding site are distinct. R16/K17 of the lectin domain is close to integrin αvβ3 and E88 of the lectin domain is close to PSGL-1 peptide and glycan. <u>**b.** Docking simulation of interaction between open/active αvβ3 (1L5G.pdb) and the lectin domain of CD62P (1g1q.pdb) was performed using Autodock3.</u> The amino acid residues selected for mutagenesis are shown. <u>c and d. The binding of the lectin domain mutants to soluble integrins αvβ3 or αIIbβ3. e. Alignment of P-, L-, and E-selectins.</u> R16/K17 and E88 are conserved in selectins. Data are shown as means +/- SD (n=3). Statistical analysis was done by ANOVA in Prism7.

It has been proposed that ligand binding to the lectin domain closes loop 83–89 around the Ca^2+^ coordination site, enabling Glu-88 to engage Ca^2+^ and fucose (13). All three selectins require Glu-88 to sustain bonds with sLex-containing ligands under force. Mutating Glu-88 to Asp (the E88D mutation) locks selectins in their functionally inactive states and markedly impairs selectin-mediated cell rolling under flow (10). To test how glycan binding and integrin binding are related, we generated the lectin domain mutants defective in integrin binding. We selected several amino acid residues within the predicted integrin-binding interface of the lectin domain for mutagenesis. We found that several mutants (R16E/K17E, K58E, K66E/K67E, K84E/R85E) were defective in binding to soluble αIIbβ3 and αvβ3 in 1 mM Mn^2+^ (Fig. 2c and 2d). Notably, the E88D mutation did not affect integrin binding, indicating that glycan binding and integrin binding sites are distinct. However, since the K84E/R85E mutation reduced integrin binding, it is likely that the glycan and integrin binding sites may be close to or overlap each other. Positions of the amino acid residues are shown in Fig. 2e.

### Inhibitors of CD62P-PSGL-1 interaction did not block the lectin domain-integrin interaction

Since the binding sites for glycan ligands and integrins are close to each other in the lectin domain, it is possible that currently available antagonists to CD62P also inhibit integrin binding. However, we found that a widely used monoclonal antibody P8G6 against CD62P did not reduce integrin binding to the lectin domain of CD62P (data not shown). This antibody has been reported to block CD62P-induced platelet aggregation (14). Also, PSGL-1-Fc fusion protein did not affect the binding of soluble integrins αvβ3 and αIIbβ3 to the lectin domain. Non-carbohydrate small-molecular weight CD62P inhibitor_KF38789 is known to block adhesion of U937 monocytic cells (PSGL-1-positive, integrin-positive) to immobilized to CD62P, but not to immobilized sLe^x^ (15). This inhibitor did not block integrin binding to the lectin domain. These findings are consistent with a model that glycans binding and integrin binding to the lectin domain are independent, and integrins and glycans may simultaneously bind to the lectin domain.

### The binding of the lectin domain to α4β1 and α5β1

CD62P is expressed on activated platelets and on activated endothelial cells and expected to support cell adhesion to endothelial cells or cancer cells by binding to αvβ3. CD62P is also expected to bind to leukocytes, but αvβ3 is not a major integrins in leukocytes. We found that the lectin domain of CD62P can interact with α4β1 and α5β1 (Fig. 4). We thus propose that CD62P binds to leukocytes through β1 integrins.

**Fig. 3.**
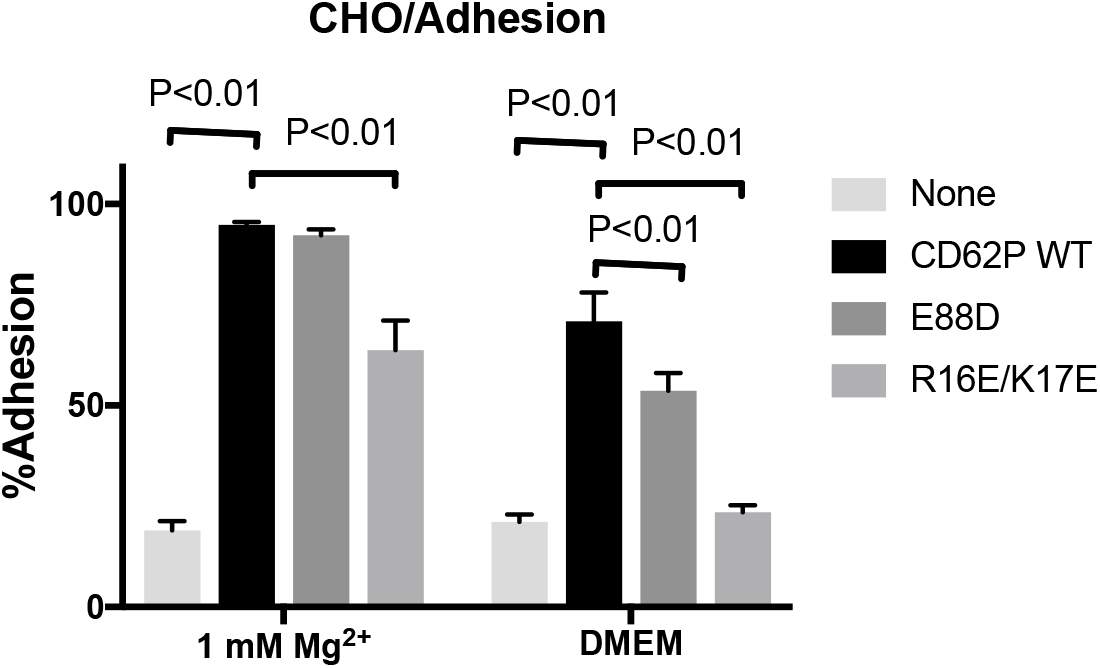
Adhesion of Chinese Hamster Ovary (CHO) cells (PSGL-1 negative) to the CD62P lectin domain. PSGL-1 is expressed in leukocytes, but not in CHO cells. Wells of 96-well microtiter plate were coated with the lectin domain (WT and mutants, coating concentration at 50 μg/ml) and remaining protein-binding sites were blocked with BSA. Wells were incubated with CHO cells (α5β1+) in Tyrode-HEPES/1 mM Mg^2+^ or DMEM. The E88D mutant is defective in binding to glycan ligand and the R16E/K17E mutant is defective in integrin binding.

**Fig. 4.**
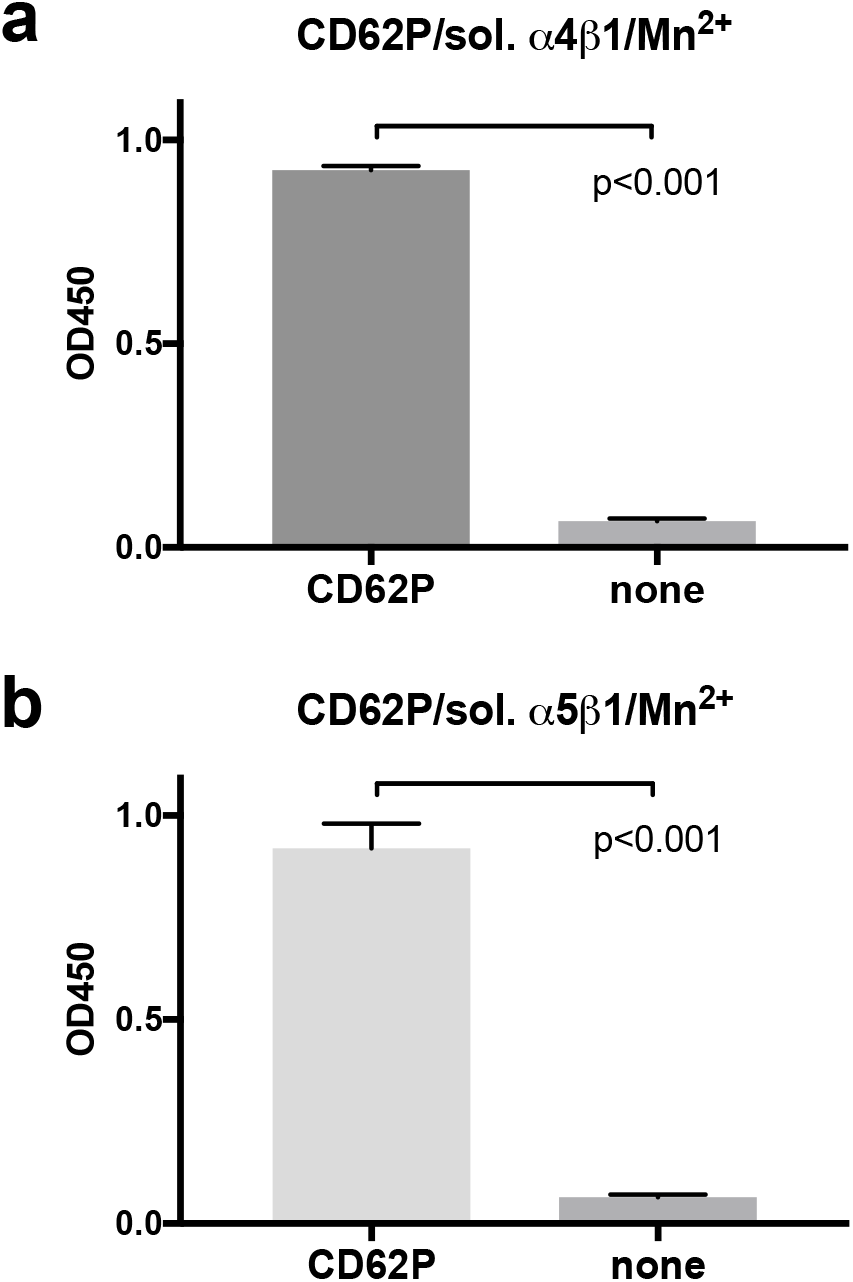
The binding of the lectin domain of CD62P to integrins α4β1 and α5β1. The binding of biotinylated α4β1 or α5β1 to immobilized CD62P lectin domain was determined in 1 mM Mn^2+^ as described in Fig. 1 except that bound integrin was quantified using streptavidine conjugated with HRP.

### The lectin domain supports static cell adhesion in a PSGL-1-independent manner

We found that immobilized WT CD62P lectin domain supported adhesion of CHO cells (α5β1+ and αvβ3 low) that do not express PSGL-1 (Fig. 3). WT CD62P supported adhesion of CHO cells (70%) in DMEM in which integrins are not activated due to high Ca^2+^ (> 1 mM), whereas WT CD62P supported more strongly (>90%) cell adhesion than in DMEM in Tyrode-HEPES buffer with 1 mM Mg^2+^, in which integrins are more activated than in DMEM. This is consistent with the idea that CD62P supports cell adhesion by binding to integrins. E88D supported cell adhesion to a level comparable to that of WT CD62P in 1 mM Mg^2+^. This is quite different from the effect of the E88D mutation on cell rolling on PSGL-1 under flow (10), indicating that CD62P-integrin interaction and CD62P-PSGL-1 interaction are distinct. The R16E/K17E mutation, which reduced integrin binding in ELISA-type binding assay, showed reduced cell adhesion in 2 mM Mg^2+^ (to 60%) and did not support cell adhesion in DMEM. Arg-16 and Lys-17 are not part of the glycan-binding region of CD62P, which is consistent with the idea that glycan binding and integrin binding are separate functions of CD62P. These findings are consistent with the model in which the lectin domain supports adhesion of CHO cells in cation-dependent and PSGL-1-independent manner.

### The lectin domain of CD62P activates soluble integrins αvβ3 and αIIbβ3 in 1 mM Ca^2+^ in a cell-free conditions

It has been proposed that CD62P primes leukocyte integrin activation during inflammation (16). However the specifics of the mechanism of priming have not been established. We have reported that several integrin ligands (e.g., fractalkine, SDF-1, sPLA2-IIA, CD40L) bound to the allosteric site of integrins (site 2) and activated integrins (8, 9, 17, 18). We hypothesized that the lectin domain of CD62P binds to site 2 and allosterically activates integrins. To test this possibility, we used ELISA-type activation assays, in which soluble integrins αvβ3 and αIIbβ3 were incubated with immobilized fibrinogen fragments, γC399tr and γC390-411 specific to αvβ3 and αIIbβ3, respectively, and incubated with soluble integrins in the presence of the lectin domain in 1 mM Ca^2+^ (to keep integrins inactive). Bound soluble integrins were determined using anti-β3 antibody. We found that the lectin domain enhanced the binding of soluble integrins to ligands (Fig. 5). This indicates that the lectin domain activated integrins. We needed high concentration of the soluble lectin domain to detect CD62P-induced integrin activation. CD62P is a transmembrane protein and soluble CD62P binds to proteoglycans on the cell surface. Therefore, CD62P is highly concentrated on the surface. Therefore, CD62P-integrin interaction is biologically relevant.

**Fig. 5.**
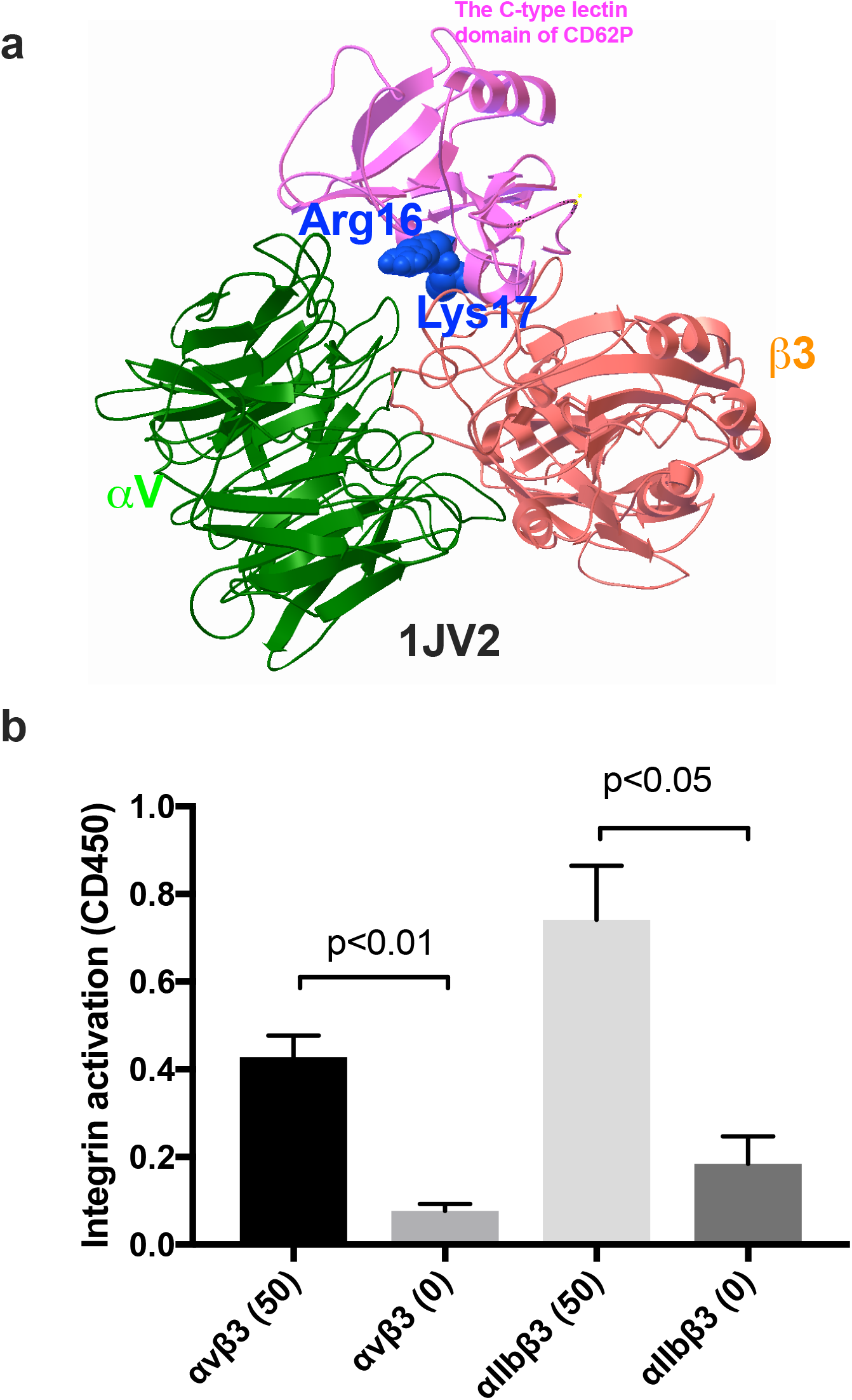
Activation of integrin αvβ3 by the lectin domain. **(a) A docking model of the lectin domain binding to site 2 of αvβ3.** Docking simulation was performed as described in the method section. **(b) Activation of soluble integrins αvβ3 and αIIbβ3 by the lectin domain in ELISA-type activation assays.** Wells of 96-well microtiter plate were coated with ligands (γC399tr for αvβ3 at 50 μg/ml, and γC390-411 for αIIbβ3 at 20 μg/ml) and the remaining protein binding sites were blocked with BSA. Wells were incubated with soluble integrins (1 μg/ml) and the lectin domain (0 or 50 μg/ml) in Tyrode-HEPES buffer with 1 mM Ca^2+^ for 1 hr, and bound integrins were quantified using anti-β3 mAbs and HRP-conjugated anti-mouse IgG.

## Discussion

The present study establishes for the first time that the C-type lectin domain of CD62P bound to integrins and support integrin-mediated cell adhesion. Notably, the present study defines the role of integrins in CD62P-mediated cell-cell interaction in the pathogenesis of diseases. Since integrins are widely expressed in different cell types, CD62P-integrin interaction may be involved in wide variety of cell-cell interaction, including several known CD62P-mediated cell-cell interaction (Fig. 6). This is in contrast to PSGL-1, which is limited to leukocytes.

**Fig. 6.**
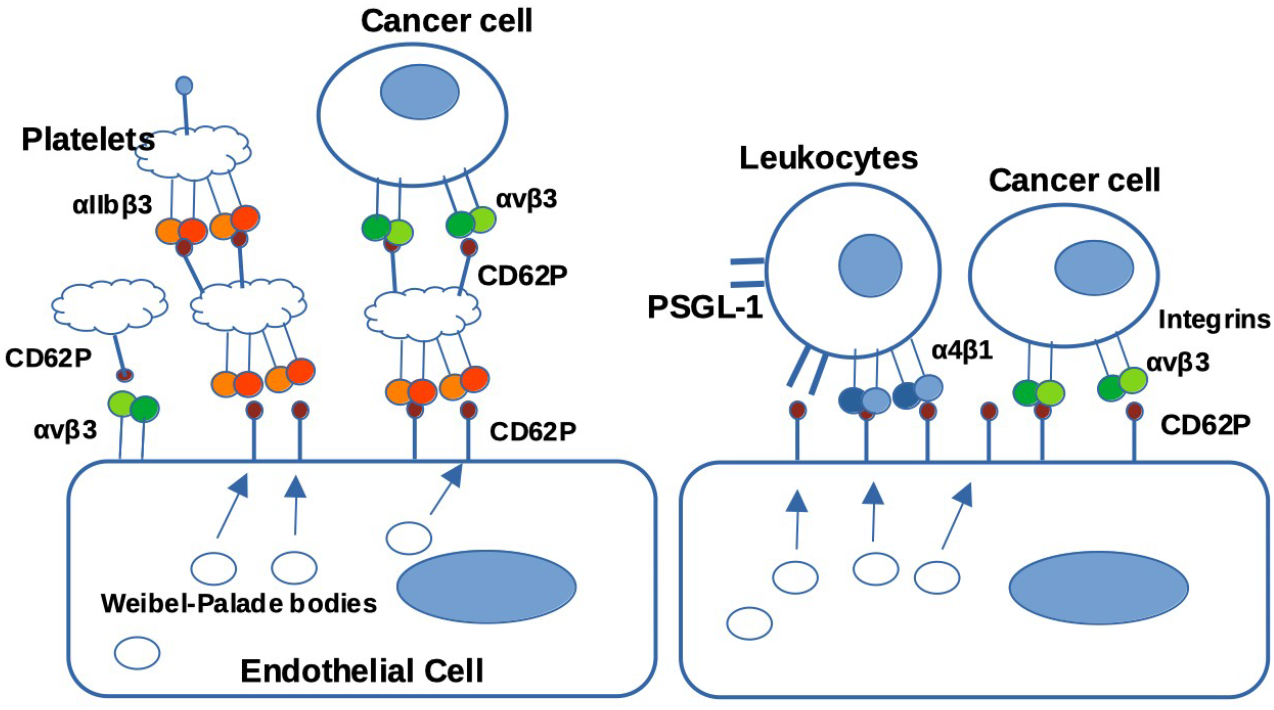
Potential role of CD62P-integrin interaction. Our findings predict that CD62P (soluble and membrane) mediates cell-cell interaction and signaling by binding to integrins. a) <u>Leukocyte rolling</u>. Leukocytes roll on the surface of endothelial cells by binding of leukocyte integrins (e.g., α4β1) to CD62P, in addition to PSGL-1 (on leukocytes) binding to CD62P. b) <u>Platelet integrin activation</u>. It is well established that αIIbβ3 is activated by inside-out signaling induced by platelet agonists. In addition to this process, our preliminary studies predict that, when platelets are activated by platelet agonists, CD62P inside the platelet is rapidly transported to the surface and activates αIIbβ3 probably by binding to the allosteric site of αIIbβ3. c) <u>Platelet-endothelial cell interaction</u>. Our preliminary studies predict that platelets bind to endothelial cells by the binding of platelet αIIbβ3 to CD62P on activated endothelial cells. d) <u>Cancer metastasis</u>. Cancer cells bind to platelets by binding of integrin αvβ3 (on cancer cells) to CD62P (on platelets). e) <u>Inflammatory signals</u>. CD62P induces signals through integrin signaling pathways, in addition to PSGL-1. This can be applied to both transmembrane and soluble PSGL-1.

### Leukocyte extravasation

It has been well established that CD62P on activated endothelial cells is involved in tethering of leukocytes by binding to PSGL-1 on leukocytes. The present study suggests that CD62P mediates cell-cell interaction by binding to integrins on leukocyte in addition to binding to glycans (PSGL-1). Also, we showed that CD62P can activate integrins by binding to site 2 in an allosteric manner. This may be critical for inducing CD62P-integrin interaction, since leukocyte integrins may not be activated in circulation.

### Platelet aggregation and integrin activation

CD62P is expressed on activated platelets and can interact with integrin αIIbβ3 on apposing platelets, leading to platelet-platelet interaction. Also, CD62P can directly activate αIIbβ3 by binding to site 2 in cis. We hypothesize that CD62P-integrin interaction and αIIbβ3 activation are key events in platelet functions.

### Metastasis

Previous studies showed that CD62P on activated endothelial cells or activated platelets is involved in tumor metastasis (19), but ligands for CD62P on cancer cells have not been fully established. Integrin αvβ3 is known to be over expressed in many cancer cells, and related to cancer metastasis. We hypothesize that CD62P-αvβ3 binding and activation of αvβ3 by CD62P in trans may be involved in interaction between cancer cells and endothelial cells, and between cancer cells and platelets.

The present study indicates that integrin-CD62P interaction is a new therapeutic target. However, we were not able to show integrin antagonists or CD62P antagonists block integrin-CD62P interaction in our hands. This may be because currently available antagonists were selected for blocking integrin binding to extracellular matrix ligands or CD62P binding to glycans.

## Experimental procedures

### Materials

Antibody P8G6 (Santa Cruz Biotechnology), KF38789 (Tocris Bioscience), and PSGL-1-Fc (Sino Biological) were obtained from the described sources.

### The C-type lectin domain and The C-type lectin and the EGF domains

The cDNA fragments encoding the C-type lectin and the C-type lectin and the EGF domains were chemically synthesized and subcloned at the BamHI/EcoRI site of pET28a. Protein expression was induced by IPTG in E. coli BL21 and purified in Ni-NTA-affinity chromatography under denaturing conditions, and refolded as described (20).

#### Binding of soluble integrins to the lectin domain

ELISA-type binding assays were performed as described previously (20). Briefly, wells of 96-well Immulon 2 microtiter plates (Dynatech Laboratories, Chantilly, VA) were coated with 100 μl PBS containing the CD62P lectin domain for 2 h at 37°C. Remaining protein binding sites were blocked by incubating with PBS/0.1% BSA for 30 min at room temperature. After washing with PBS, soluble recombinant αIIbβ3 (AgroBio, 1 μg/ml) was added to the wells and incubated in HEPES-Tyrodes buffer (10 mM HEPES, 150 mM NaCl, 12 mM NaHCO3, 0.4 mM NaH2PO4, 2.5 mM KCl, 0.1% glucose, 0.1% BSA) with 1 mM MnCl_2_ for 1 h at room temperature. After unbound αIIbβ3 was removed by rinsing the wells with binding buffer, bound αIIbβ3 was measured using anti-integrin β3 mAb (AV-10) followed by HRP-conjugated goat anti-mouse IgG and peroxidase substrates.

#### Activation of soluble integrins by the lectin domain

ELISA-type binding assays were performed as described previously (17). Briefly, wells of 96-well Immulon 2 microtiter plates were coated with 100 μl PBS containing γC399tr (for αvβ3) and γC390-411 (for αIIbβ3) for 2 h at 37°C. Remaining protein binding sites were blocked by incubating with PBS/0.1% BSA for 30 min at room temperature. After washing with PBS, soluble recombinant αIIbβ3 (AgroBio, 1 μg/ml) was pre-incubated with the lectin domain for 10 min at room temperature and was added to the wells and incubated in HEPES-Tyrodes buffer with 1 mM CaCl2 for 1 h at room temperature. After unbound integrins was removed by rinsing the wells with binding buffer, bound integrins was measured using anti-integrin β3 mAb (AV-10) followed by HRP-conjugated goat anti-mouse IgG and peroxidase substrates.

#### Docking simulation

Docking simulation of interaction between CD62P (Protein Data Bank code 1G1Q), and integrin αvβ3 was performed using AutoDock3, as described (21). In the current study, we used the headpiece (residues 1–438 of αIIb and residues 55–432 of β3) of αvβ3 (Protein Data Bank code 1L5G, open headed). Cations were not present in αIIbβ3 during docking simulation (22, 23). The classical ligand-binding site (site 1) or the allosteric site (site 2) of αvβ3 was selected as a target for the lectin domain. To perform docking simulation of the interaction between site 2 of closed headed αvβ3, we used 1JV2.pdb.

#### Statistical analysis

Treatment differences were tested using ANOVA and a Tukey multiple comparison test to control the global type I error using Prism 7 (GraphPad Software).

**Table 1.** Amino acid residues in interaction between CD62P and αvβ3 (1L5G.pdb) predicted by docking simulation.

| CD62P | $\alpha\text{v}$ | $\beta 3$ |
| --- | --- | --- |
| Asn13, Ile14, **Arg16**, **Lys17**, Tyr18, Gln20, Asn21, Thr24, Asp25, **Lys55**, Asn56, Lys58, Thr61, Val63, Gly64, Thr65, Lys66, Lys67, Ala68, Asn83, Lys 84, Arg85, Asn86, Asn87, Asp89, His108, Leu110, Ala120, Cys122, Gln123, Asp124, Met125, Lys129, Glu132, Cys133, Glu135, Tyr145 | Ala149, Asp150, Phe177, Tyr178, Gln180, Arg211, Thr212, Ala213, Gln214, Ala215, Ile216, Asp218, Arg218, | Tyr122, Ser123, Met124, Lys125, Asp126, Asp127, Trp129, Ser130, Gln132, Asn133, Lys137, Asp179, Met180, Lys181, Thr182, Arg214, Asp251, Ser334, Met335, Asp336, Ser337, Asn339, Val340, Leu341, Gln342 |

**Table 2.** Amino acid residues in interaction between CD62P and αvβ3 (1JV2.pdb) predicted by docking simulation.

| CD62P | $\alpha\text{v}$ | $\beta 3$ |
| --- | --- | --- |
| Thr7, Lys8, Trp12, Asn13, Ile14, **Arg16**, **Lys17**, Tyr18, Gln20, Asn21, Arg22, Tyr23, Thr24, Asp25, Arg54, Lys55, Asn56, Asn57, Lys58, Thr59, Val63, Gly64, Thr65, Lys66, Lys67, Ala68, Asn86, Asn87, Asp89, Leu110, Tyr118, Thr119, Ala120, Ser121, Cys122, Gln123, Asp124, Met125, Glu135 | Glu15, Asn44, Thr45, Thr46, Pro48, Gly49, Ile50, Val51, Glu52, Gly76, Asn77, Asp79, Asp83, Asp84, Pro85, Phe88, His91, Gln120, Arg122 | Lys159, Pro160, Val161, Met165, Ile167, Ser168, Glu171, Ala172, Glu174, Asn175, Pro176, Cys177, Tyr178, Asp179, Met180, Lys181, Thr183, Cys184, Pro186, Met187, Phe188, **Val275**, **Gly276**, **Ser277**, **Asp278**, **His280**, **Tyr281**, **Ser282**, **Ser284**, **Thr285**, **Thr286**, |

## Author contributions

Y. K. T. performed the experiments. Y. T. conceived the project, designed the experiments, analyzed the data, and wrote the article.

## Funding and additional information

This project was supported by Pilot funding from Comprehensive Cancer Center at UC Davis School of Medicine. This work is partly supported by the UC Davis Comprehensive Cancer Center Support Grant (CCSG) awarded by the National Cancer Institute (NCI P30CA093373).

